# Rubisco activase remodels plant Rubisco via the large subunit N-terminus

**DOI:** 10.1101/2020.06.14.151407

**Authors:** Jediael Ng, Oliver Mueller-Cajar

**Affiliations:** School of Biological Sciences, Nanyang Technological University, 60 Nanyang Drive, Singapore 637551. Singapore

**Keywords:** Rubisco, Rubisco activase, ATPases associated with diverse cellular activities (AAA), photosynthesis, molecular chaperone, carbon fixation

## Abstract

The photosynthetic CO_2_ fixing enzyme ribulose 1,5-bisphosphate carboxylase/oxygenase (Rubisco) forms inhibited complexes with multiple sugar phosphates, including its substrate ribulose 1,5-bisphosphate. At least three classes of ATPases associated with diverse cellular activities (AAA+ proteins) termed Rubisco activases (Rcas) have evolved to remodel inhibited Rubisco complexes. The mechanism of green-type Rca found in higher plants has proved elusive, because until recently higher plant Rubiscos could not be expressed recombinantly. Towards identifying interaction sites between Rubisco and Rca, here we produce and characterize a suite of 33 Arabidopsis Rubisco mutants for their ability to be activated by Rca. We find that Rca activity is highly sensitive to truncations and mutations in the conserved N-terminus of the Rubisco large subunit. Both T5A and T7A substitutions cannot be activated by Rca, but present with increased carboxylation velocities. Our results are consistent with a model where Rca functions by transiently threading the Rubisco large subunit N-terminus through the axial pore of the AAA+ hexamer.

## INTRODUCTION

Virtually all carbon dioxide that enters the biological world does so via the Calvin Benson Bassham cycle (1). Aerobic autotrophic organisms such as plants, algae and cyanobacteria all utilize this somewhat suboptimal CO_2_-fixation process, which is dependent on catalysis by the slow and promiscuous enzyme Rubisco. The enzyme binds the five carbon sugar Ribulose 1,5-bisphosphate (RuBP), adds a carbon dioxide molecule and hydrolyzes the six carbon intermediate to form two molecules of 3-phosphoglycerate (3PGA). In plants each active site only processes ∼1-3 reactions per second, and frequently oxygen gas is incorporated instead of CO_2_, which leads to production of the toxic metabolite 2-phosphoglycolate.

To overcome these flux limitations, Rubisco is overexpressed to constitute up to 50% of the leaf soluble protein and is believed to be the most abundant protein on earth (2,3). Recognition that the enzyme catalyzes the rate limiting step have made its performance the heartpiece of multiple ongoing crop improvement strategies (4,5). In addition to its slow speed and inaccuracy, the enzyme is also susceptible to form dead-end inhibited complexes with several sugar phosphates that are present in its environment (6,7). CO_2_ fixation ceases, unless the inhibitors are constantly removed. This action is performed by a group of dedicated molecular chaperones that have been termed the Rubisco activases (Rcas) (8-10). Three classes of Rca exist, and although all belong to the superfamily of AAA+ proteins, their primary sequences and mechanisms are highly distinct, indicating convergent evolution(11,12). Red-type Rca found in red-lineage phytoplankton and proteobacteria transiently threads the C-terminus of the Rubisco large subunit through the axial pore of the AAA+ hexamer (13-15). In contrast the CbbQO-type Rca found in chemoautotrophic proteobacteria consists of a cup-shaped AAA+ hexamer (CbbQ_6_) bound to a single adaptor protein CbbO, which is essential for Rubisco activation (9). During Rca function the hexamer remodels CbbO, which is bound to inhibited Rubisco via a von Willebrand Factor A domain(16).

The detailed molecular mechanism by which inhibitory compounds are removed by higher plant Rca-mediated modelling of Rubisco’s active site has long remained elusive (11,17). As functional Rca could be produced recombinantly, a large volume of biochemical information has accumulated on Rca variants (18,19). In summary the data supports a canonical AAA+ pore-loop threading mechanism where the flat top surface of the hexameric disc engages Rubisco, followed by axial pore-loop threading of an element of Rubisco(20,21). The intrinsically disordered N-terminal domain, especially a conserved tryptophan, is also important in engaging the holoenzyme (21,22). Regarding Rubisco, early studies using green algal *Chlamydomonas* Rubisco were able to pinpoint two residues on the Rubisco large subunits’ βC-βD loop that contact the specificity helix (H9) of Rca (23,24). However, a historical inability to produce plant Rubisco in heterologous organisms such as *Escherichia coli* hampered further progress. This hurdle was recently removed (25), and here we took advantage of the newly established capability to produce and biochemically characterize many plant Rubisco variants for their interaction with Rca. We find that the highly conserved RbcL N-terminus is essential for Rca function, with a particular importance of two threonine residues T5 and T7. This is consistent with an N-terminal pore loop threading mechanism for higher plant Rca.

## RESULTS

### A surface scan of higher plant Rubisco for Rca-interacting residues

We used the recently established *E. coli* plant Rubisco expression platform (25,26) to produce a series of Arabidopsis Rubisco large subunits variants mutated in surface-localized residues in an effort to discover additional regions important for protein-protein interactions. We first tested the βC-βD loop mutations E94K and P89A as positive controls (Fig. 1*A*), as these substitutions had earlier been shown to greatly perturb the ability of Spinach Rca to activate Chlamydomonas Rubisco (27). We then assayed the fully activated holoenzyme (ECM) and the inhibited apo enzyme bound to the substrate ribulose 1,5-bisphosphate (ER) in the presence and absence of the short (Rcaβ) isoform of Arabidopsis Rca (Fig. 1*B*). Consistent with the Chlamydomonas-spinach result, the inhibited E94K variant of Arabidopsis Rubisco remained non-functional in the presence of its cognate Rca, reconfirming the importance of the N-terminal βC-βD loop in the interaction. The P89A variant, however, was activated well in this system (Fig. 1*C*), suggesting that the βC-βD loop – Rca interaction is less sensitive to mutation when using Arabidopsis proteins.

**Figure 1.**
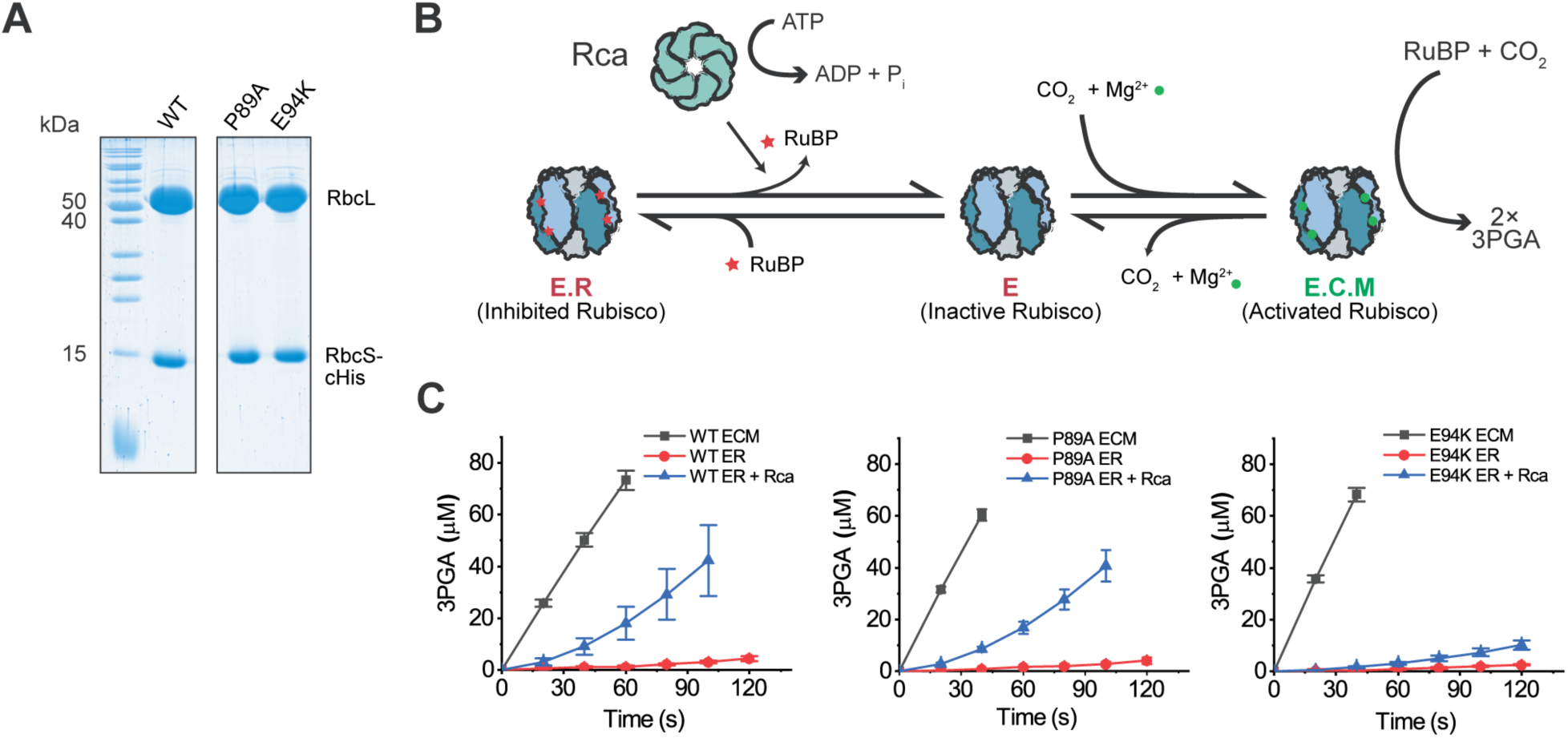
Production and Rca assay of plant Rubisco variants. (*A*) SDS-PAGE analysis of purified recombinant wild-type Arabidopsis Rubisco and βC-βD mutant variants (*B*) Rca functions to remodel inhibited E.R. complexes, releasing RuBP. The apo-enzyme E can then bind non-substrate CO_2_ and Mg^2+^ to form the functional E.C.M. holoenzyme. (*C*) Rubisco activation assays of wild type and βC-βD loop mutant Rubisco variants. Assays were performed at 25°C and contained 0.5 µM Rubisco active sites in the presence and absence of Arabidopsis Rca (2 µM protomer). Error bars indicate the S.D. of at least three independent experiments.

We next targeted a range of surface-localized Rubisco large subunit residues for mutagenesis (Fig. 2*A*, Fig. S2). As we have found earlier, multiple positively charged residues on the face of the Rca disc are important for its ability to activate Rubisco (21,28), and therefore the chosen mutations were biased towards probing negatively charged surface residues. This included those located in a negatively charged pocket at the dimer-dimer interface that has recently been implicated in the binding of carboxysomal Rubisco linker proteins in prokaryotic green-type Rubiscos (Fig. 2*B*)(29,30). We successfully purified 17 variants (Fig. S1), which were all able to carboxylate RuBP similarly to wild-type (Fig. 2*C*, Fig. S3). Rca assays indicated that the different variants could still be activated effectively indicating the chosen residues were not of critical importance to the Rubisco-Rca interaction (Fig. 2*C*, Fig. S3). Only K14A showed a statistically significant 52% increase in its Rca-mediated activation rate, possibly reflecting a reduced stability of the inhibited complex. Clearly the chosen single amino-acid substitutions were insufficient to disrupt the extensive protein-protein interaction interface involved in Rubisco activation. However, attempts to produce combinations of mutations were unsuccessful either due to insolubility or non-functionality for all tested cases.

**Figure 2.**
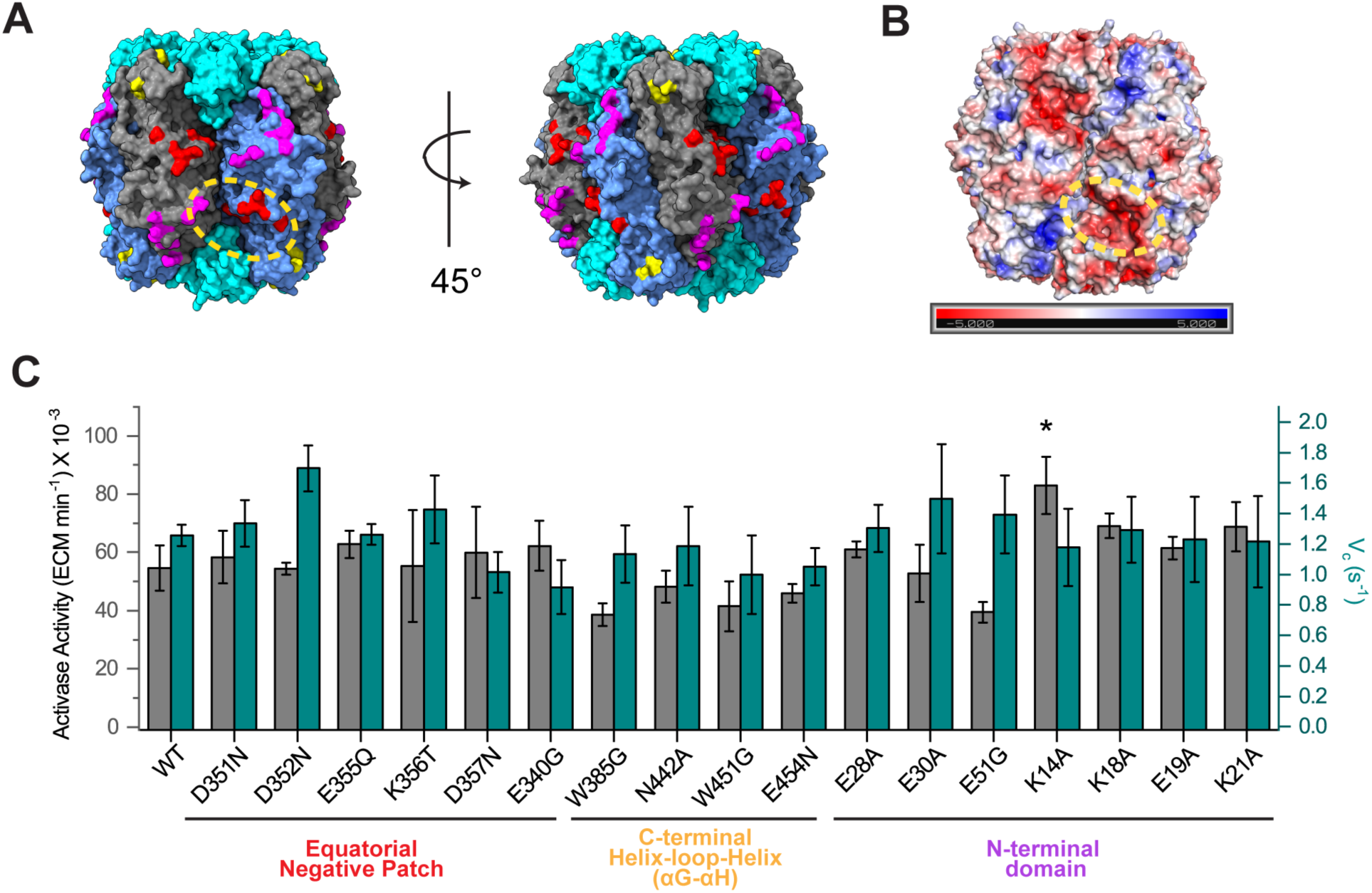
Mutational scan of the Rubisco surface. (*A*) Mutated surface residues are visualized on the surface of inhibited Arabidopsis rubisco (PDB: 5IUO) and coloured in groups. (*B*) Electrostatic surface potential as analysed by the Adaptive Poisson-Boltzmann Solver (APBS) wrapper function in PyMOL. The location of a negatively charged patch between two large subunit dimers is circled in yellow. (*C*) Rca function and carboxylation rates of surface mutants in groups corresponding to (*A*). Assays were conducted at 25°C using 0.5 µM Rubisco active sites (ER or ECM) and 2 µM wild type activase protomers when appropriate. Error bars indicate the S.D. of at least three independent experiments. The corresponding time course data is shown in Fig. S2. A significant difference from the wild-type value is indicated by an asterisk (One-way ANOVA, posthoc Tukey test, p < 0.05).

### The RbcL N-terminus is essential for Rca function

The red-type Rubisco activase CbbX transiently threads the RbcL C-terminus (13-15). However, the C-terminus of green-type Rubisco large subunits is poorly conserved and is of variable length (31), indicating a distinct mechanism for green-type Rca function. In contrast, while sequences at the N-terminus of red-type Rubisco large subunits differ between species, both length and sequence of the N-terminus of higher-plant RbcL is essentially completely conserved (Fig. 3*A*). In available crystal structures, residues 8-20 of the N-terminus are ordered only when the active site is in the closed (ligand-bound) form (Fig. 3*B*). In the closed conformation, the N-terminus is positioned directly above the 60’s loop that co-ordinates P1 of the substrate, with F13, K14, G16 and K18 forming interactions with multiple residues of the 60’s loop (32). Coupled with evidence that residues 9-15 of Rubisco from wheat are essential for functional carboxylation activity (33), the stringent conservation of the first 8 residues suggested thus a tantalizing target for mutational analysis.

**Figure 3.**
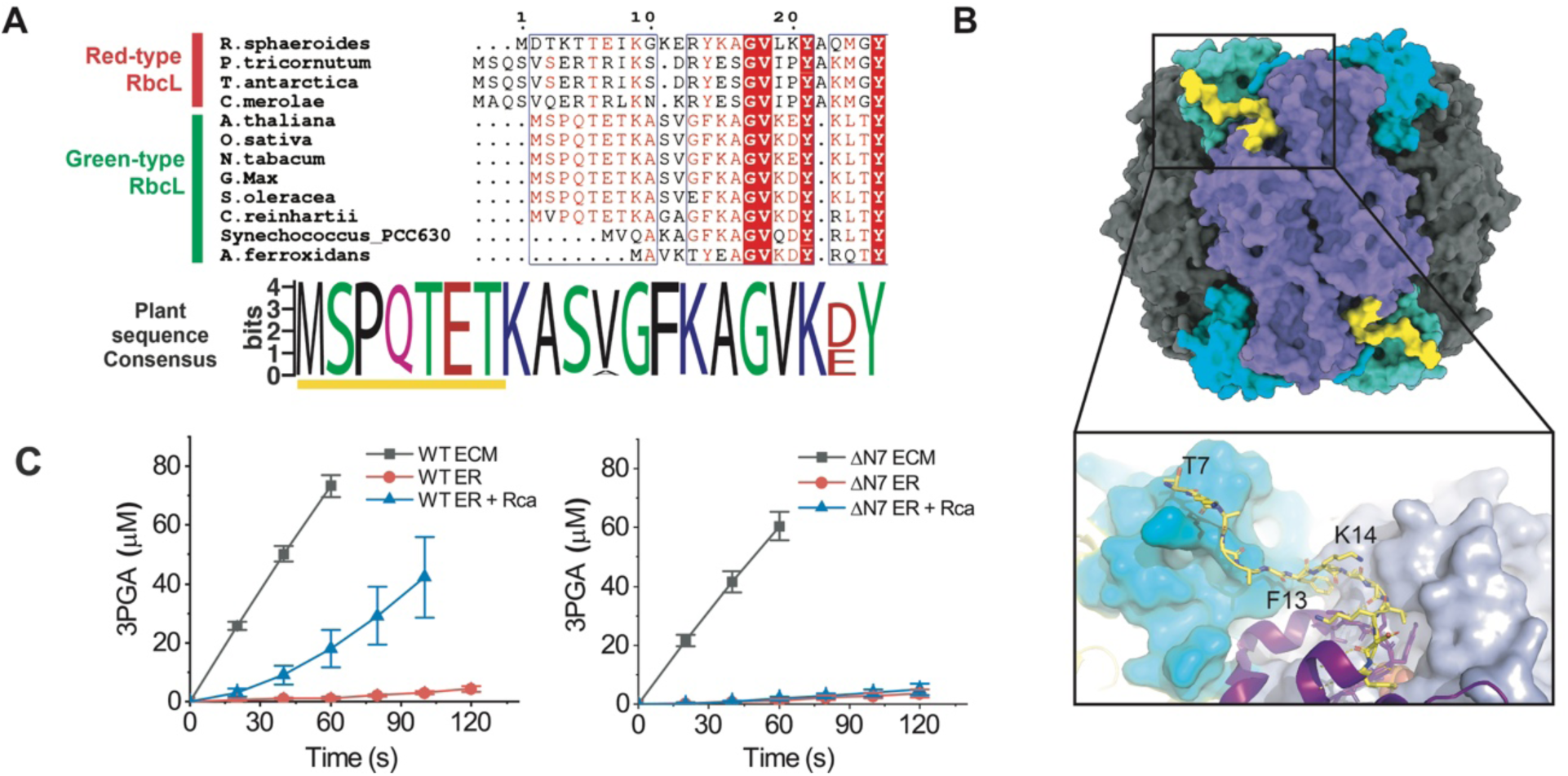
The N-terminus of the Rubisco large subunit is essential for Rca function. (*A*) Sequence alignment of RbcL N-terminal sequences. In higher plants the N-termini are almost completely conserved, as indicated by a web-logo representation of the consensus sequence (∼500 plant RbcL sequences). (*B*) Visualization of the ordered N-terminal segment (yellow-starting from residue 7) using a model of the inhibited *Chlamydomonas reinhardtii* Rubisco (PDB:1GK8). (*C*) Rubisco activase assays of wild-type and ΔN7 mutant Arabidopsis Rubiscos. Assays were conducted at 25°C with 2µM wild type activase protomers and 0.5 µM Rubisco active sites. Error bars indicate the S.D. of at least three independent experiments.

A Rubisco variant with the first seven amino-acids replaced by methionine (ΔN7) displayed 83 % of wild-type carboxylation velocity (Fig. 3*C*). However, when the ER complex was formed, Rca was unable to reactivate ΔN7 (Fig. 3*C*). This result was consistent with the notion that a higher plant Rca hexamer engages the disordered N-terminus via its axial pore loops, followed by transient threading leading to active site disruption and inhibitor release.

### A dissection of the RbcL N-terminal binding motif

We then performed a detailed mutational analysis of the RbcL N-terminus, generating a series of variants that, in the ECM form were all able to carboxylate at least as well as wild-type (Fig. 4). Shortening the N-terminus by one or two amino acids (ΔN1, ΔN2) did not negatively affect Rca function. In contrast, replacing the first four amino acids with methionine (ΔN3) or deleting residues 5-7 (ΔTET) almost completely eliminated the ability of Rca to activate Rubisco (Fig 3A).

**Figure 4.**
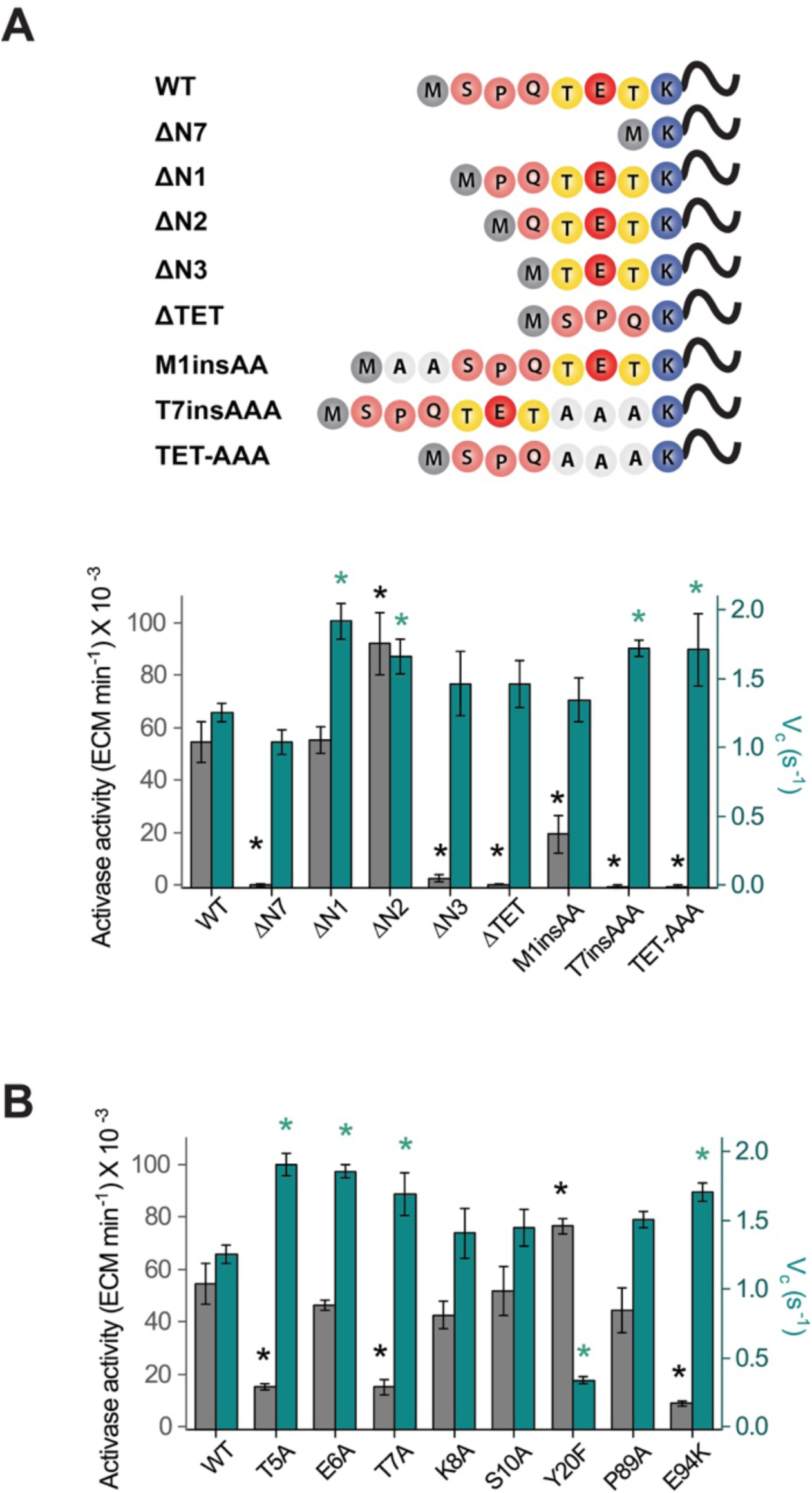
Mutational analysis of the RbcL N-terminus. Activase activity and carboxylation rates of Rubisco N-terminal truncations and insertions (*A*) and point mutant (*B*) variants were assayed. Time courses are shown in Fig. S4. Values significantly different from their wild-type equivalent are indicated by an asterisk (One-way ANOVA, posthoc Tukey test, p < 0.05).

Lengthening the N-terminus by inserting two alanine residues upstream of residue 2 (M1insAA) greatly reduced Rca function by ∼64 %. Changing the register of the N-terminal sequence by inserting a AAA sequence upstream of K8 (T7insAAA) in the wild-type or ΔTET variant (TET-AAA) also eliminated function. These results indicated that Rca function was highly sensitive to both length and identity of the N-terminus.

Next, we evaluated the effect of single amino acid substitutions in the N-terminal motif. Whereas E6A and K8A substitutions were well tolerated, both T5A and T7A resulted in ∼70% reductions in Rca functionality. This findings indicates that the two threonine residues are likely to play an important role in the threading process, possibly via specific interactions with residues in Rca’s pore-loop 1 and 2 (20,21). We also further note that the observed 2 amino acid step interval would be consistent with successive zipper-like interactions that have been described to be utilized for substrate engagement at the central pore by multiple other AAA+ proteins (34-38).

Y20 forms a hydrogen bond to E60, a key catalytic residue that interacts with K334, which is positioned at the apex of Loop 6, and thought to orient the CO_2_ molecule for gas addition (39). We hypothesized that this interaction could act to disrupt the active site, when the N-terminus is displaced by Rca-threading. The Y20F Rubisco mutant had a ∼73% reduced carboxylation rate, but the activation rate by Rca was increased by 39% (Fig. 4*B*). This result suggests that the Y20-E60 interaction is important for the integrity of the active site, and its loss facilitates disruption of the inhibited complex.

Quite unexpectedly, we found that multiple N-terminal variants and E94K presented with significantly enhanced carboxylation velocities (up to 53%) compared to the wild-type enzyme under the conditions used in our spectrophotometric Rubisco assay (Fig. 4A,B). This suggests that reducing the interactions of the N-terminus with the enzyme may result in faster Rubiscos. Interestingly, the fast cyanobacterial Form IB enzyme from Synechococcus PCC6301 (40) possesses a truncated N-terminus (equivalent to ΔN6) (Fig. 3*A*). In a follow-up study, it will be important to use ^14^C-CO_2_ fixation assays to accurately quantify the carboxylation kinetics (41) and the CO_2_/O_2_ specificity factor of the fast N-terminal variants.

## DISCUSSION

The availability of *E. coli* produced recombinant higher plant Rubiscos permitted us to rapidly produce many variants and assay their capability to be engaged and activated by their cognate Rca chaperone. Mutational analysis of the holoenzyme surface indicated that Rca compatibility was not easily disrupted, with the tested variants remaining functional (Fig. 2). Arguably, inclusion of less conservative substitutions such as charge switches, could have been more informative here.

In contrast, mutagenesis of the highly conserved N-terminus resulted in multiple variants that were able to carboxylate RuBP, but could not be activated by Rca. The best described conserved mechanism of numerous AAA+ ATPases concerns the translocation of a substrate peptide through the central pore of the hexamer (42), and this is the modus operandi of the red-type Rca (13). Green-type Rca pore-loops have been shown to be critical for Rubisco activation (20,21). In addition we have long been aware of the RbcL βC-βD loop – Rca specificity helix H9 interaction (27,43,44). Assuming an axial pore loop- RbcL N-terminal threading mechanism we can now further constrain the positioning of an Rca hexameric model (20) in relation to an inhibited Rubisco structure (45). Helix 9 elements of two adjacent Rca subunits can be placed in proximity to two RbcL βC-βD loops that are located on two adjacent dimers (Fig. 5). In this configuration the N-terminal tail (missing 6 amino-acids in the structure used) is then accessible to the Rca pore. Transient threading would then result in pulling residues 13 to 20 away from the large subunit body. Disruptions of the associated van der Waal’s and polar interactions (32), especially with the RbcL 60’s loop, may be sufficient to trigger active site opening and inhibitor release.

**Figure 5.**
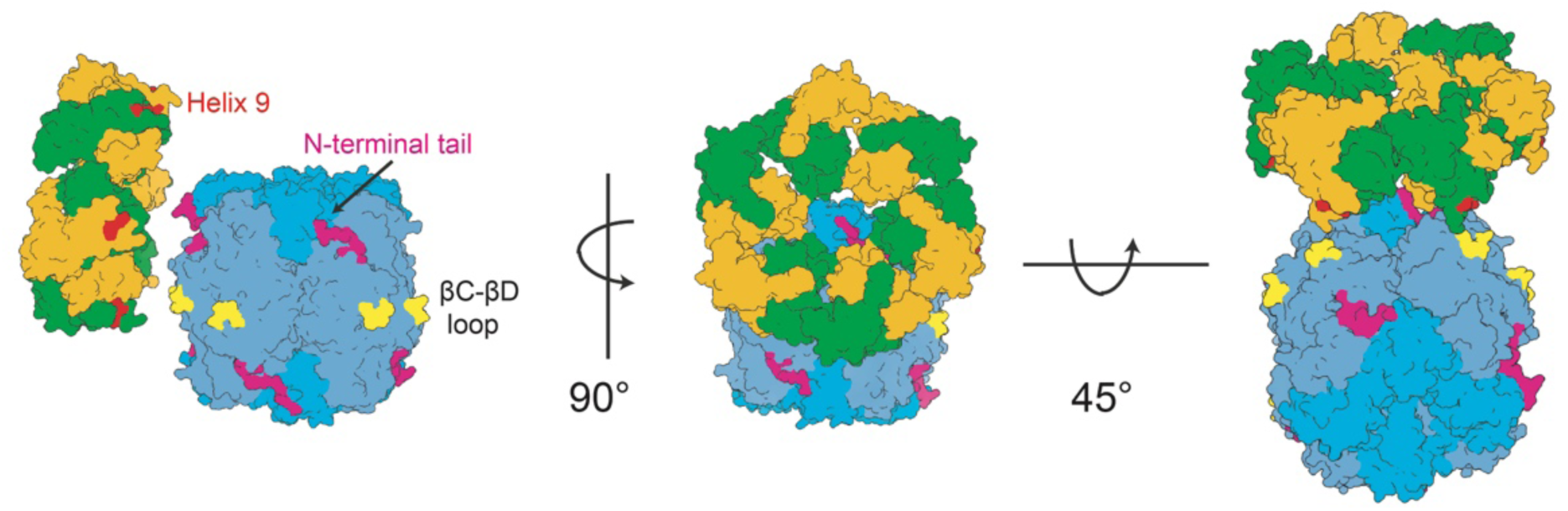
Model for the Rubisco-Rca interaction. By placing the Helix 9 interaction site (in red) of two adjacent Rca subunits in proximity to the βC-βD loops (yellow) of two large subunits belonging to different dimers, the RbcL N-terminus (in magenta-N-terminal 6 amino acids missing) can be positioned under the axial pore of the Rca hexamer. Transient pore-loop threading would then lead to disruptions of interactions between the N-terminus and the catalytic 60s loop, followed by inhibitor release. PDB:1GK8 (Rubisco) and 3ZW6 (Rca).

An important question that remains completely unaddressed is the role of the critical Rca N-terminal domain. This disordered stretch of ∼60 amino acids is not resolved in structural models, and a single amino acid substitution of W15A eliminates Rca function (21,22). It is likely involved in an additional, so far undescribed, anchoring site.

The model is consistent with an exquisite structural snapshot of a prokaryotic carboxysome-associated green-type Rca hexamer bound to cyanobacterial Rubisco that has been communicated in a concurrent Biorxiv pre-print (46). In agreement with our findings, the study also reports that an N-terminal 9-amino acid truncation of the tobacco Rubisco large subunit abolishes tobacco Rca function. Green-type Rubisco activation is thus an ancient, conserved process that appears to precede the primary endosymbiotic event (11).

## MATERIALS AND METHODS

### Molecular biology

Plasmids pBAD33k-*At*RbcLS, p11a-*At*C60αβ/C20 and pCDFduet-*At*R1/R2/Rx/B2 enabling the production of Arabidopsis Rubisco in *E. coli* were a gift from Dr. Manajit Hayer-Hartl (25). To achieve our final construct containing the large and small subunits of Rubisco, a 6x Histidine tag was appended to the C-terminus of rbcS via the Quikchange protocol (Stratagene). Restriction free cloning of the RBS-*At*RbcLScHis cassette was utilized to insert the cassette into the multiple cloning site 1 of the pRSFDuet™-1 plasmid (Novagen). To obtain single mutants, the Quikchange protocol was applied to pRSFduet-*At*RbcLScHis. Truncations of the N-terminus were performed by PCR amplification of regions flanking the unwanted sequence. Linearized products were then phosphorylated by T4 PNK (NEB) before end to end ligation. All primers used are listed in Table S1 and protein-encoding sequences were verified by DNA sequencing.

To obtain a vector encoding *Arabidopsis thaliana* Rca (AtRcaβ), the sequence corresponding to amino acid residues 59 to 474 (Uniprot P10896) were amplified from a cDNA library of Arabidopsis with BamHI and NotI restriction sites at the 5’ and 3’ end respectively. The sequence was then inserted into the multiple cloning site of the pHue expression vector using the appropriate restriction sites to yield the final construct pHueAthRcaβ.

### Protein Purification

Recombinant wild-type activase from Arabidopsis were expressed and purified following our protocol for the purification of *Agave tequilana* activases to yield activases with a single glycine prior to the native N-terminus of the enzyme (28). For expression and purification of recombinant Rubiscos, *E.coli* BL21 cells containing p11a-*At*C60αβ/C20, pCDFduet-*At*R1/R2/Rx/B2, and pRSFduet-*At*RbcLScHis were grown in LB medium supplemented with ampicillin (200 µg mL^-1^), kanamycin (30 µg mL^-1^), and streptomycin (50 µg mL^-1^). Starter cultures of 2 mL were grown overnight at 37°C prior to inoculation of 1 L cultures. Large scale cultures were grown for 3 hours at 37°C to reach an OD600 of 0.3 - 0.4 before temperatures were lowered to 23°C and harvested after 16 hours. Cells were lysed in Histrap-buffer A (50 mM Tris-HCl pH 8.0, 50 mM NaCl, 10 mM Imidazole). Soluble fractions were then applied to HisTrap HP 5 mL columns (Sigma Aldrich) equilibrated with Histrap-buffer A. Proteins were eluted with a linear imidazole gradient from 10 mM to 200 mM. Following elution, Rubisco containing fractions were immediately subjected to a Superdex 200 gel filtration column (GE Healthcare) equilibrated with buffer A (20 mM Tris-HCl pH 8.0, 50 mM NaCl) supplemented with 5% (v/v) glycerol. Pure Rubisco fractions were then pooled, concentrated, and flash-frozen for storage at −80°C. Pure proteins were quantified my measuring their absorbance at 280 nm, using extinction coefficients calculated using the ProtParam tool (https://web.expasy.org/protparam/).

### Biochemical assays

Rubisco and Rubisco reactivation activities were measured and quantified as described (14), using the spectrophotometric Rubisco assay (47). RuBP was synthesized enzymatically from ribose 5-phosphate (48) and purified by anion exchange chromatography(49). ECM was formed by incubating 20 µM Rubisco active sites in Buffer A supplemented with 20 mM NaHCO_3_ and 10 mM MgCl_2_ (50 mins, 25 °C). For ER, complexes were generated by incubating 20 µM Rubisco active sites in Buffer A containing 4 mM EDTA (10 mins) prior to addition of RuBP to a final concentration of 1 mM (50 mins, 25 °C). Rca activities were calculated using the ECM and ER carboxylase time-courses collected on the same day. All assays were performed in assay buffer (100 mM Tricine pH 8.0, 5 mM MgCl_2_) containing 3 µl coupling enzymes mixture (Creatine P-kinase (2.5 U/ml), Glyceraldehyde-3-P dehydrogenase (2.5 units/ml), 3-phosphoglycerate kinase and Triose-P isomerase/Glycerol-3-P dehydrogenase (20/2 units/ml)), 20 mM NaHCO_3_, 0.5 mM NADH, 2 mM ATP, 10 mM Creatine-P, 1 mM RuBP. A final concentration of 0.5 µM Rubisco active sites and 2 µM Rca protomer were utilized where appropriate.

## ACKNOWLEDGEMENTS

We thank Dr. Manajit Hayer-Hartl for the gift of plasmids pBAD33k-*At*RbcLS, p11a-*At*C60αβ/C20 and pCDFduet-*At*R1/R2/Rx/B2. Lynette Liew synthesized RuBP. pHue*Ath*Rcaβ was cloned by Devendra Shivhare. This work was funded by the Ministry of Education (MOE) of Singapore Tier 2 grant to O.M.-C (MOE2016-T2-2-088).

## Supporting Information (SI)

**Fig. S1.**
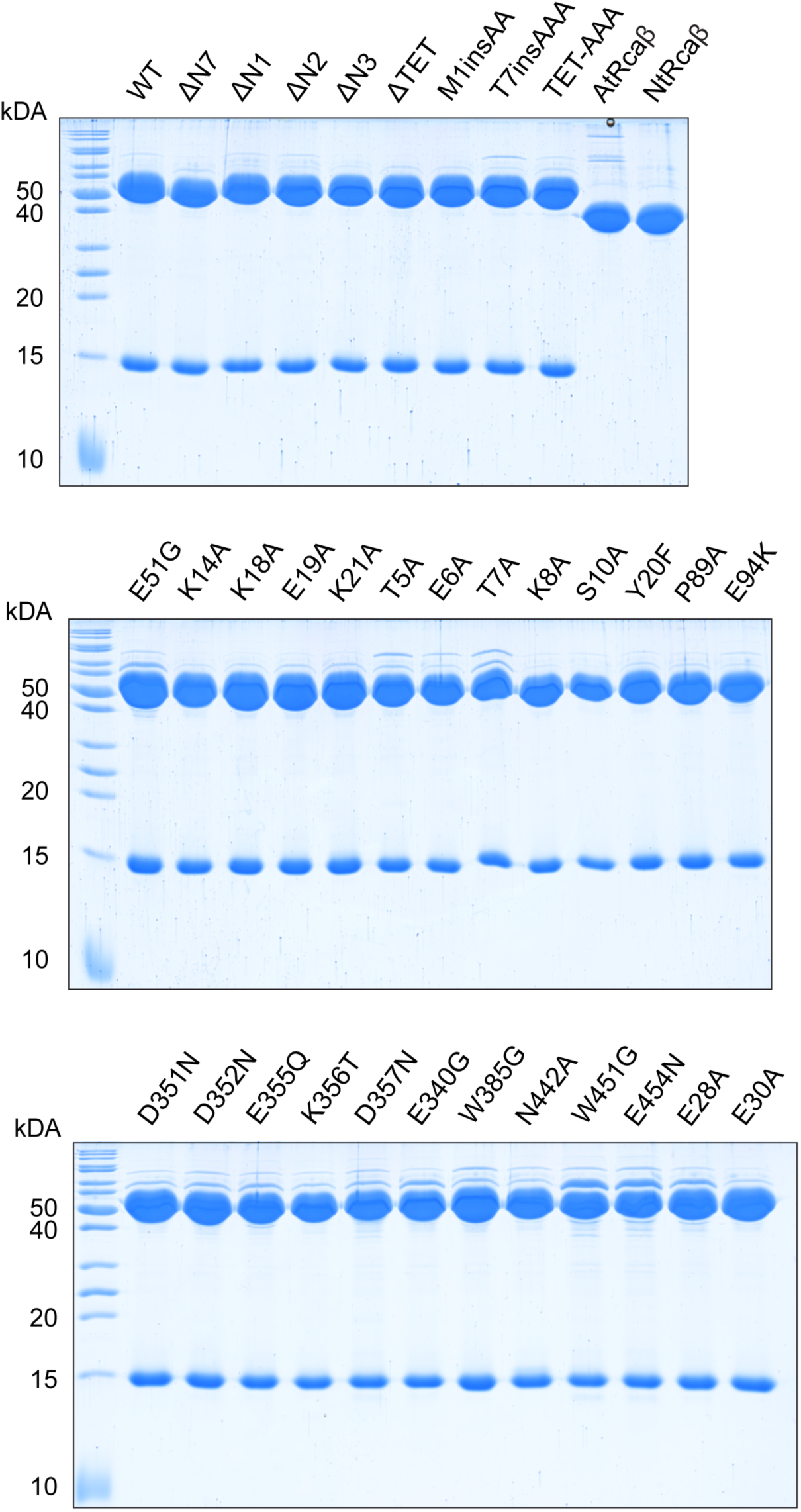
SDS-PAGE analysis of proteins used in this study. 4 µg of purified protein was loaded per lane.

**Fig. S2.**
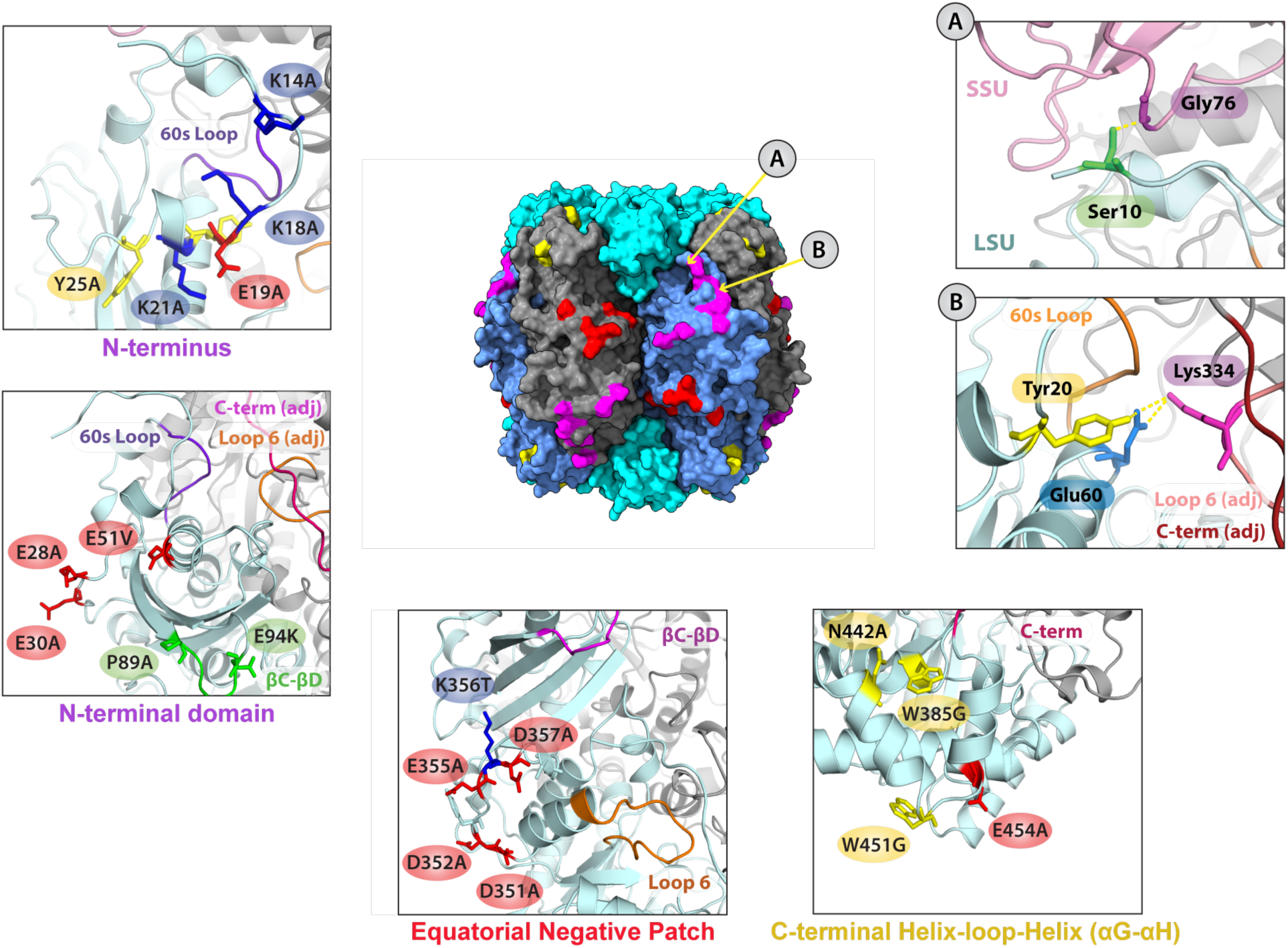
Structural representations of mutated residues on the surface of Rubisco. Detailed views of surface residues substituted in the mutational analysis. Panels on the right depict the interaction network for S10 and Y20. Serine 10 hydrogen bonds to Glycine 67 of the small subunit while the hydroxyl group on the side chain of Tyrosine 20 hydrogen bonds to Glutamate 60. Mutants of these residues were generated to investigate whether these interactions were essential for the mechanism of the activase (Fig. 4*B*).

**Fig. S3.**
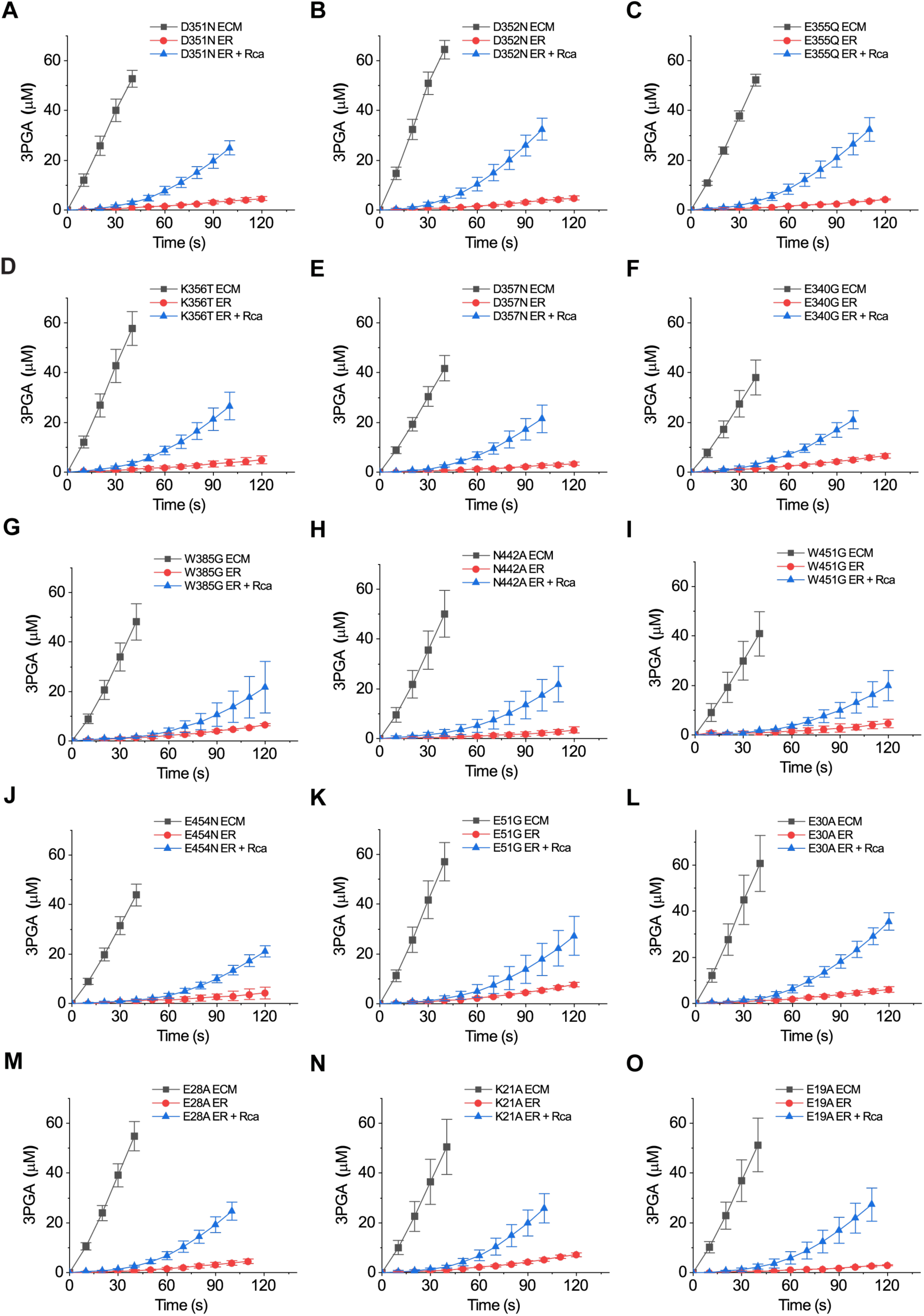
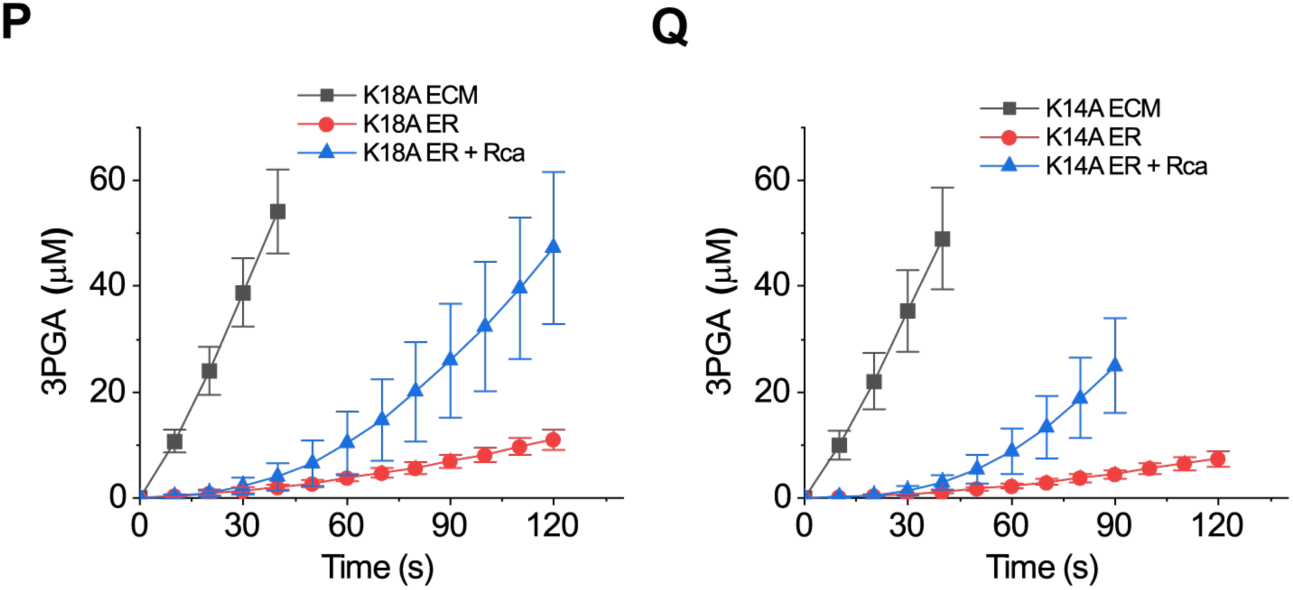
Rubisco activase assays of surface mutant variants. Carboxylation time course data of activated Rubisco (ECM), inhibited Rubisco (ER), and inhibited Rubisco in the presence of wild type Arabidopsis activase (*At*Rcaβ) is shown for all mutants described in Fig. 2C.

**Fig. S4.**
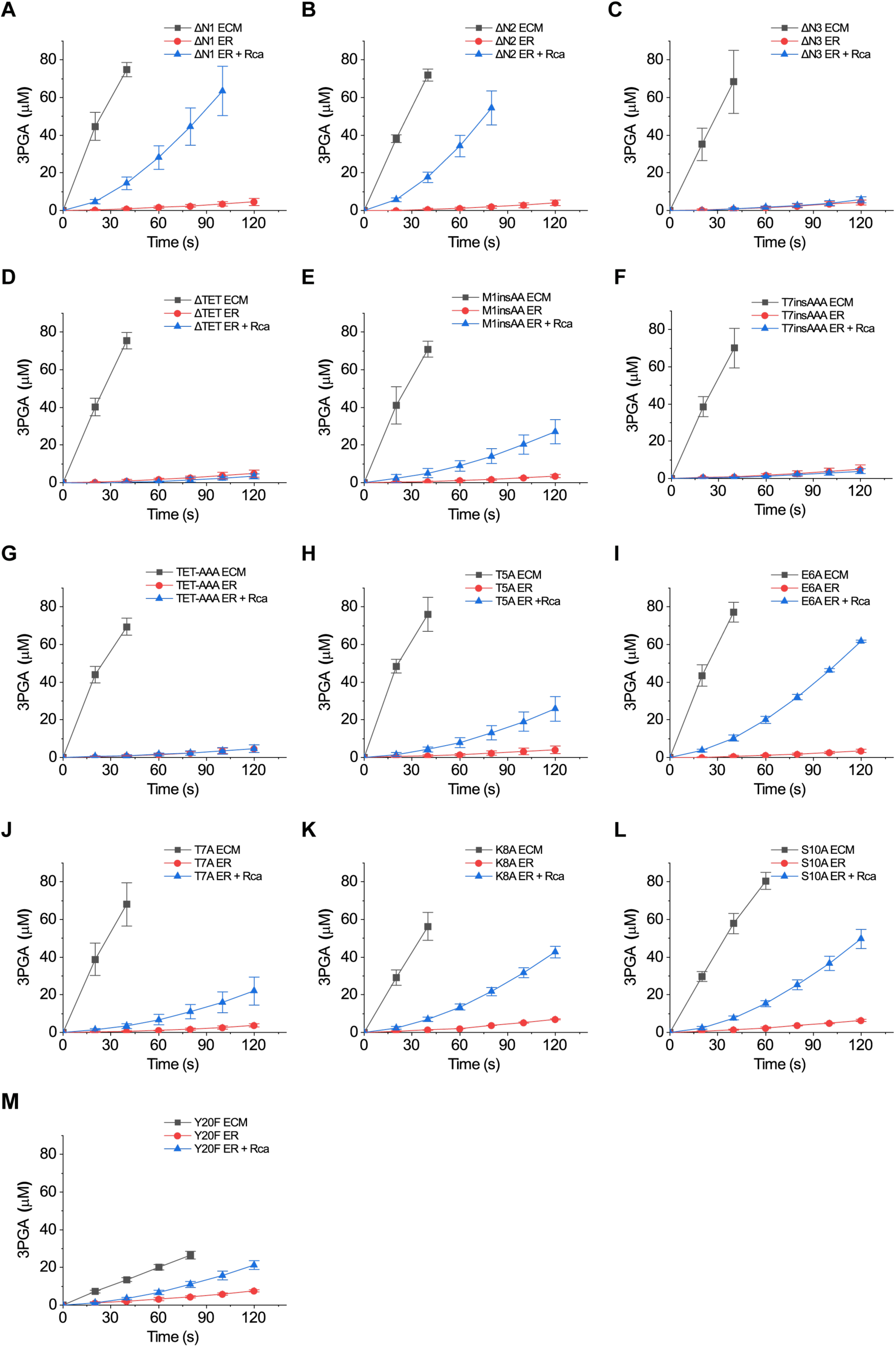
Rubisco activase assays of N-terminal mutant variants. Carboxylation time course data of activated Rubisco (ECM), inhibited Rubisco (ER), and inhibited Rubisco in the presence of wild type Arabidopsis activase (*At*Rcaβ) are shown for all mutants described in Fig. 4*A*-*B*.

**Table S1.**
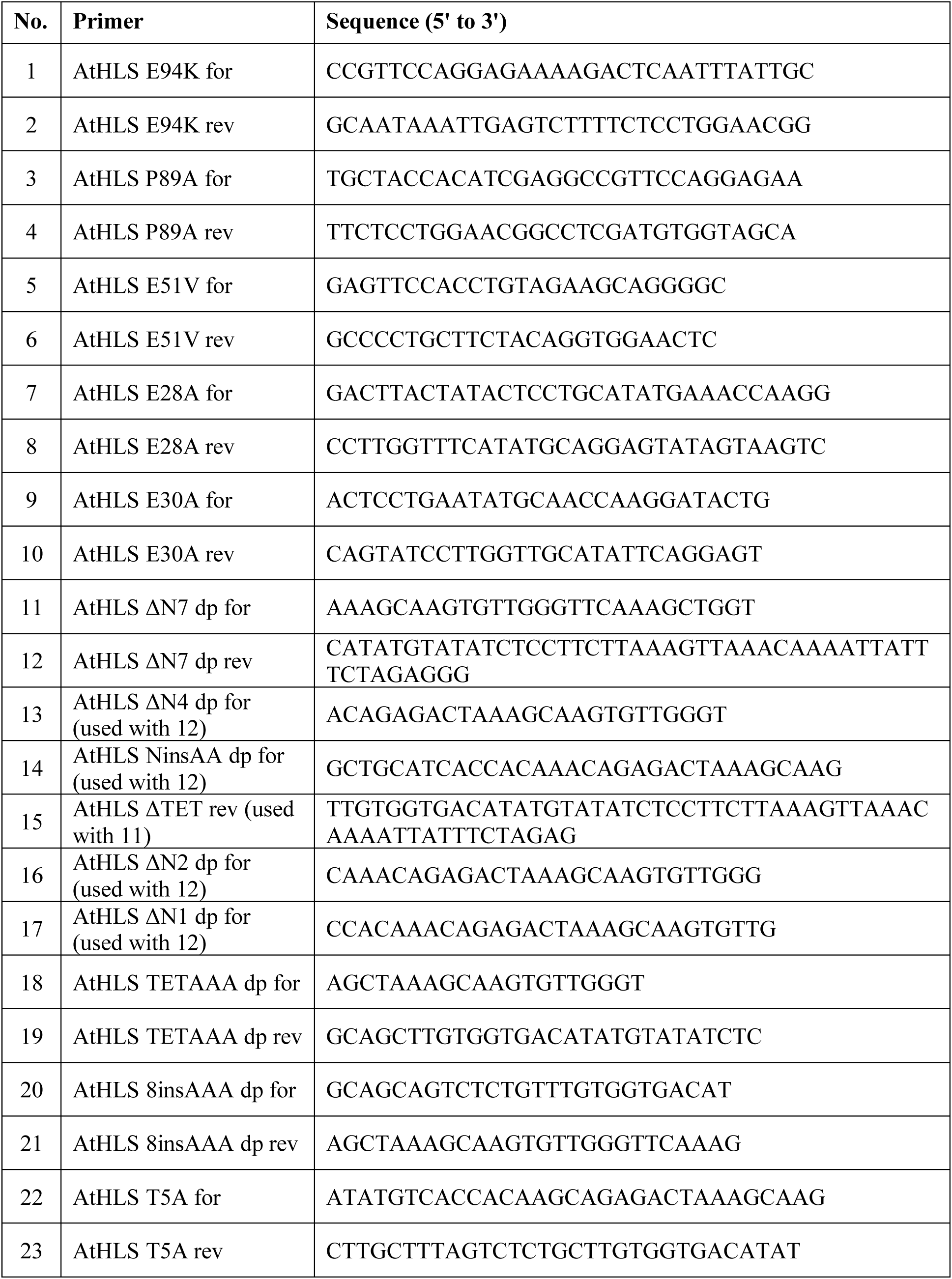
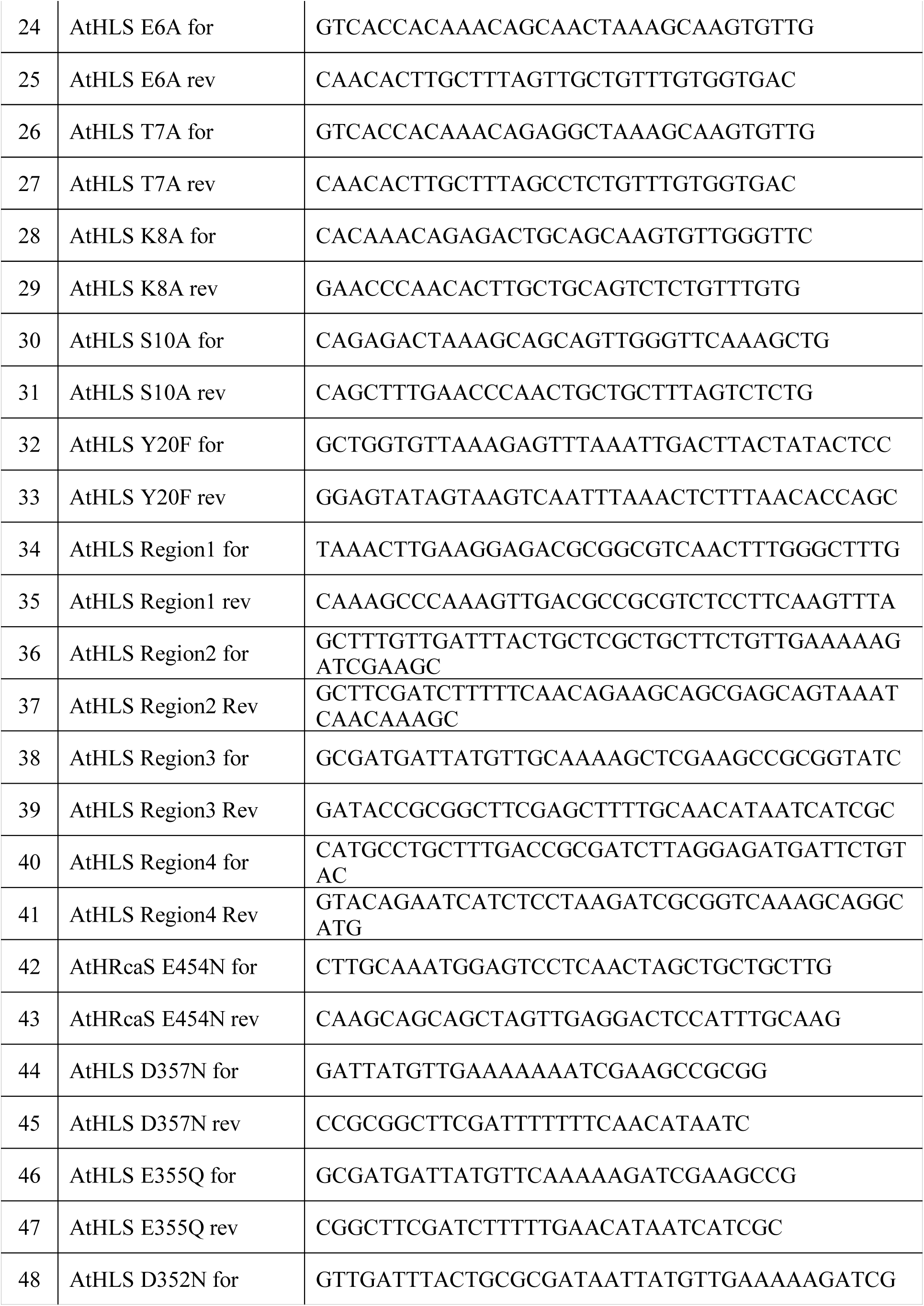
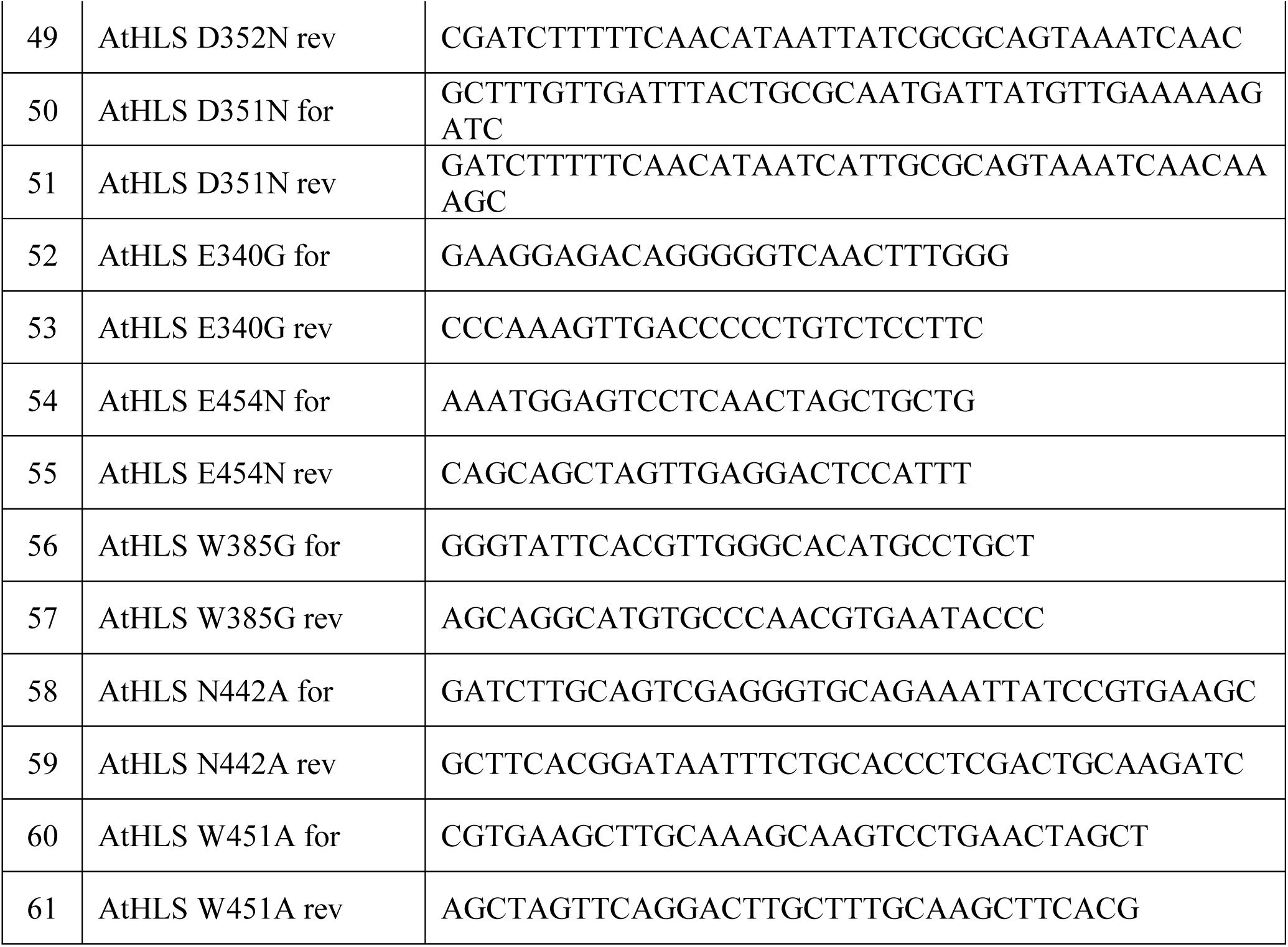
Primers used in this study

